# Comparative transcriptomics reveal distinct patterns of gene expression conservation through vertebrate embryogenesis

**DOI:** 10.1101/840801

**Authors:** Megan E. Chan, Pranav S. Bhamidipati, Heather J. Goldsby, Arend Hintze, Hans A. Hofmann, Rebecca L. Young

## Abstract

Despite life’s diversity, studies of variation across animals often remind us of our shared evolutionary past. Abundant genome sequencing over the last ~25 years reveals remarkable conservation of genes and recent analyses of gene regulatory networks illustrate that not only genes but entire pathways are conserved, reused, and elaborated in the evolution of diversity. Predating these discoveries, 19^th^-century embryologists observed that though morphology at birth varies tremendously, certain stages of embryogenesis appear remarkably similar across vertebrates. Specifically, while early and late stages are variable across species, anatomy of mid-stages embryos (the ‘phylotypic’ stage) is conserved. This model of vertebrate development and diversification has found mixed support in recent analyses comparing gene expression across species possibly owing to differences across studies in species, embryonic stages, and gene sets compared. Here we perform a comparative analysis using 186 microarray and RNA-seq expression data sets covering embryogenesis in six vertebrate species spanning ~420 million years of evolution. We use an unbiased clustering approach to group stages of embryogenesis by transcriptomic similarity and ask whether gene expression similarity of clustered embryonic stages deviates from the null hypothesis of no relationship between timing and diversification. We use a phylogenetic comparative approach to characterize expression conservation pattern (i.e., early conservation, hourglass, inverse hourglass, late conservation, or no relationship) of each gene at each evolutionary node. Across vertebrates, we find an enrichment of genes exhibiting early conservation, hourglass, late conservation patterns and a large depletion of gene exhibiting no distinguishable pattern of conservation in both microarray and RNA-seq data sets. Enrichment of genes showing patterned conservation through embryogenesis indicates diversification of embryogenesis may be temporally constrained. However, the circumstances (e.g., gene groups, evolutionary nodes, species) under which each pattern emerges remain unknown and require both broad evolutionary sampling and systematic examination of embryogenesis across species.

## Introduction

During embryogenesis, a single-cell zygote develops into a multicellular, functional embryo. Given the complexity of this process and the astonishing diversity of resultant phenotypes, the similarities in the developmental processes and anatomy of embryogenesis across species are striking and have captivated the imagination of biologists for nearly two centuries (von Baer, 1828). For example, vertebrates establish a highly conserved body plan (‘*bauplan*’) from which species-specific variation and elaborations develop. Yet, whether there are generalizable rules that direct diversification of embryogenesis across distantly-related species remains controversial (Richardson et al., 1997; Bininda-Emonds et al., 2003). Inspired by von Baer’s (1828) pioneering observations, one hypothesis suggests that early and late phases of embryogenesis are variable across species (owing to diversity and species specificity of reproductive modes and post-body plan elaboration, respectively), while anatomy mid-embryogenesis is conserved (Fig. 1). According to this ‘developmental hourglass’ hypothesis (Elinson, 1987), similarity of the mid-embryogenesis ‘phylotypic stage’ (at the pharyngula stage: Ballard, 1976, 1981; Sander, 1983; Richardson, 1995) reflects developmental constraints of body plan formation, including global signaling interdependence and interactions (Raff, 1996; Galis and Metz, 2001) and temporal and spatial patterns of Hox expression (Duboule, 1994). Still others have hypothesized that development of later stages is dependent on early stages of embryogenesis and that this ‘developmental burden’ results in highest conservation early in embryogenesis (Riedl, 1978; discussed in Irie and Kuratani, 2014) (Fig. 1).

**Figure 1.**
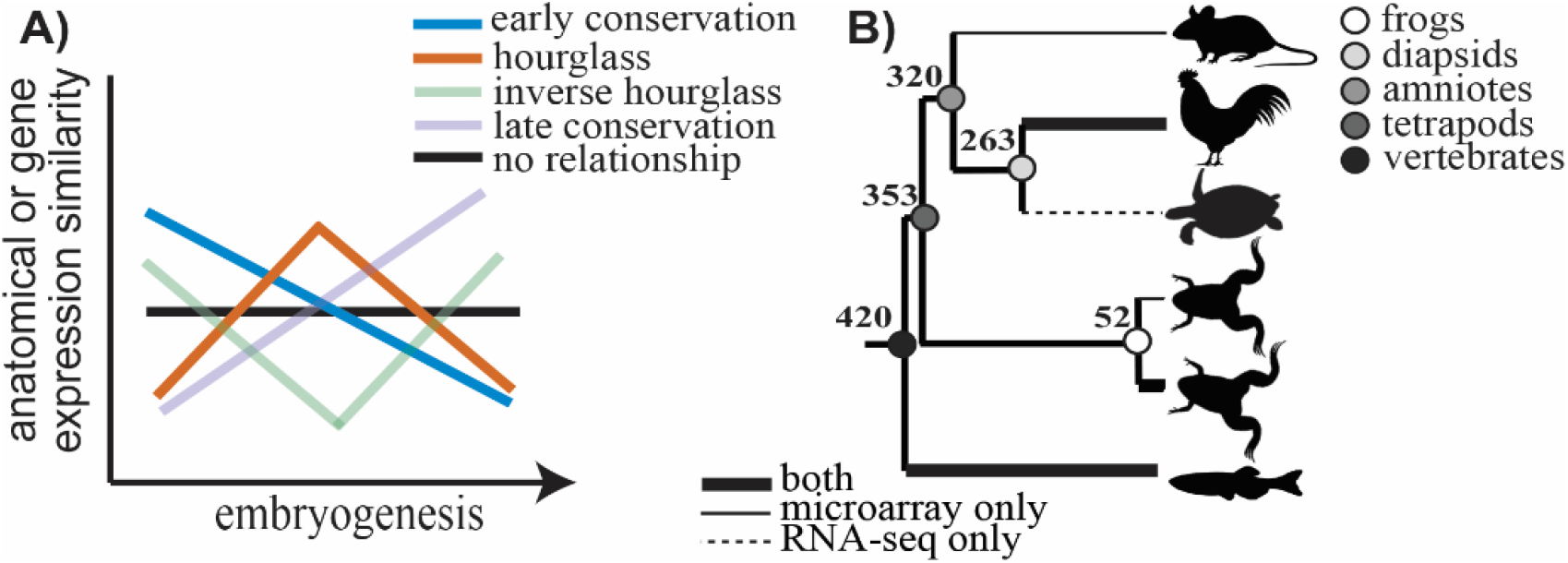
Anatomical and gene expression similarity predicted under different models of conservation through embryogenesis (A). Expression conservation was assessed using 186 publicly available microarray and RNA-seq data sets through embryogenesis across a phylogeny of six vertebrate species (B). Divergence times at each node are shown in millions of years.

Over the past decade, an increasing number of studies have leveraged the ever growing genome-scale data and approaches to test omics-level predictions of the hourglass hypothesis and its underlying mechanistic basis (Fig. 2A; Supplementary Fig. S1; Supplementary Table 1). Nevertheless, support for the hourglass model of development varies across studies and the question remains very much unsettled (Fig 2B). A number of studies comparing gene expression variation through embryogenesis across species have found support for an increase in expression conservation mid-embryogenesis (Domazet-Loso and Tautz, 2010; Irie and Kuratani, 2011, 2014; Yanai et al., 2011; Levin et al., 2012; Wang et al., 2013; Gerstein et al., 2014; Zalts and Yanai, 2017). Surprisingly, one study comparing expression divergence of animals from different phyla reported an inverse hourglass – where expression differences were highest mid-embryogenesis (Levin et al., 2016)– although this analysis did not account for phylogenetic non-independence (Dunn et al., 2018). Still others found that diversification in gene expression is not consistent with the hourglass model (Piasecka et al., 2013; Tian et al., 2013; Wu et al., 2019). von Baer’s (1828) original morphological observations were based on vertebrate embryos separated by ~420 million years of evolution; however, recent analyses varied in the evolutionary distances among species investigated (Fig. 2C), sometimes even spanning more than one phylum (de Mendoza et al., 2013; Levin et al., 2016; Hu et al., 2017) and often testing predictions of the hourglass hypothesis in only one or two species (e.g., zebrafish, Domazet-Loso and Tautz, 2010; soft-shell turtle and chicken, Wang et al., 2013; Caenorhabditis elegans, Zalts and Yanai, 2017; Fig. 2D). Taken together, even though numerous studies have used sophisticated –omics level analyses to examine embryogenesis across diverse species, we still lack conclusive molecular evidence of a developmental hourglass.

**Figure 2.**
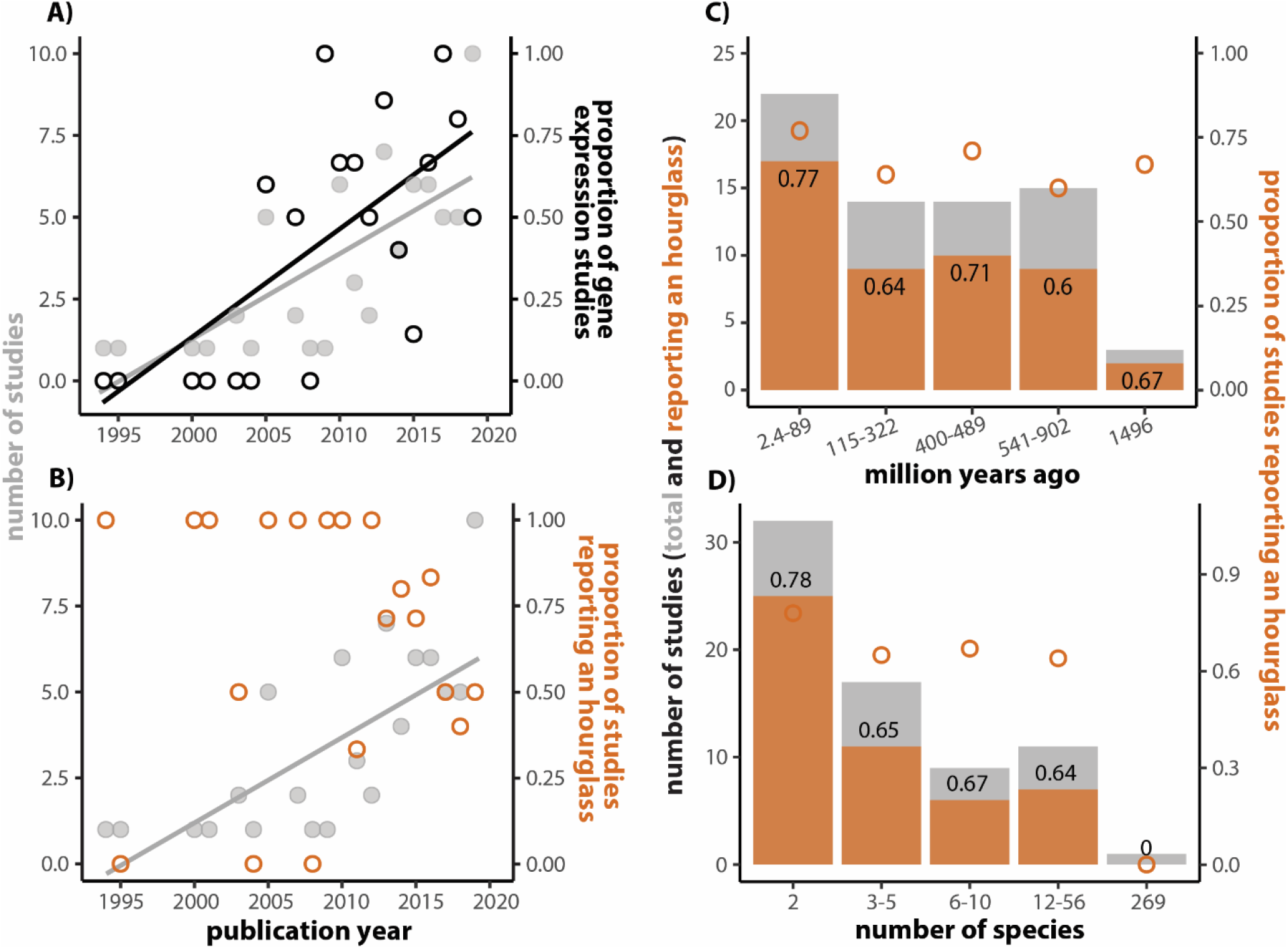
Studies testing the developmental hourglass and alternative hypotheses have increased considerably since the start of the new millennium (A, grey line: r^2^ = 0.44, F(1, 18) = 15.7, p = 0.0009, mainly driven by an increase in the number of studies focused on gene expression patterns (A, black line: r^2^ = 0.6, F(1, 18) = 29.4, p = 3.729e-05). Despite this increased attention to this question, whether variation in embryogenesis follows an hourglass pattern has remained unresolved (B, orange points: r^2^ = −0.06, F(1, 18) = 0.004, p = 0.95;). Neither divergence time of the species compared (C), nor number of species included in any given study (D) affect whether an hourglass pattern is observed. Quantitative literature analysis of studies examining early embryonic development across species was carried out according to the PRISMA flow diagram (Supplementary Fig. 1; Moher et al., 2009). Detailed methods are provided in Supplementary Materials (Supplementary Table 1, Supplementary Fig. S1)

Any test of the developmental hourglass hypothesis faces several fundamental challenges. First, proper alignment of stages of embryogenesis across species is difficult. Despite several valiant attempts to overcome this difficulty (Bininda-Emonds et al., 2003; Mungall et al., 2012; Gerstein et al., 2014; Li et al., 2014), nomenclature and sampling conventions that often vary substantially across different model systems as well as widespread heterochrony of developmental events have prevented a satisfactory solution. Irie and Kuratani (2011) circumvented this challenge by directly comparing only a subset of well-defined stages across species, but it is unclear to what extent any observed temporal pattern might be depend on the stages selected for comparison. Second, what constitutes an appropriate gene set to compare across species is very much unclear. Most studies to date have examined conservation of all expressed gene for which orthologs can be identified for all the taxa in the analysis. However, others have suggested that abundant expression of housekeeping genes may bias discovery of gene expression conservation across species (Piasecka et al., 2013). Third, it is clear that the genomic and developmental processes underlying even anatomically similar and homologous phenotypes can diverge via developmental drift or selective processes (de Beer, 1971; Wagner, 1989; True and Haag, 2001; Wilkins, 2002; McGary et al., 2010; Young and Wagner, 2011). As a result, some authors have argued that studies examining the evolution of organismal phenotypes should focus on a core set of regulatory genes critical to the initiation of the specific developmental program of that character (designated as “kernels” by Davidson and Erwin, 2006, or “Character Identity Networks, ChINs” by Wagner, 2007). However, how to identify the relevant gene set (i.e., kernel or ChIN) that is fundamental for shared developmental processes of vertebrate embryogenesis is unclear. Finally, it is often not clear what the appropriate null expectation should be in comparative studies, as it may depend on the type of data available and the level of analysis (Dunn et al., 2018; Young and Hofmann, 2019; Church and Extavour, 2020).

While numerous studies have reported patterned expression divergence (i.e., early conservation or hourglass patterns) through embryogenesis across species, what pattern is followed and whether that diversification pattern varies over evolutionary time remains unclear. Here, we perform a comparative analysis using 186 publicly available microarray and RNA-seq expression data sets covering embryogenesis in six vertebrate species spanning ~420 million years of evolution. We use an unbiased clustering approach to group stages of embryogenesis by transcriptomic similarity and ask whether gene expression similarity of clustered embryonic stages deviates from the null hypothesis that gene expression levels are invariant of developmental time. Second, using a phylogenetic comparative analysis, we characterized the expression conservation pattern (i.e., early conservation, hourglass, inverse hourglass, late conservation, or no relationship) exhibited by each gene at each evolutionary node and ask whether the number of genes that fall into a given pattern deviates from a biologically meaningful null expectation. Finally, we discuss challenges of comparative analyses that rely on publicly available transcriptome data and suggest several novel approaches for future tests of the hourglass hypothesis.

## Methods

### Obtaining and preprocessing genome-wide gene expression data from public repositories

Gene expression profiles through embryogenesis were obtained from publicly available repositories for six vertebrate species. In total 112 microarray data sets from five species and 74 RNA-seq data sets from four species were included in the analysis. Developmental time points included for each species and gene expression profiling technology are detailed in Supplementary Tables 2 and 3 (microarray and RNA-seq, respectively). Data sets include: zebrafish, *Danio rerio* (microarray: Roux and Robinson-Rechavi, 2008; RNA-seq: Comte et al., 2010; Yang et al., 2013); chicken, *Gallus gallus* (microarray: Irie and Kuratani, 2011; RNA-seq: Wang et al., 2013), Chinese soft-shell turtle, *Pelodiscus sinensis* (RNA-seq: Wang et al., 2013), mouse, *Mus musculus* (microarray: Irie and Kuratani, 2011; Xue et al., 2013); African clawed frog, *Xenopus laevis* (microarray: Yanai et al., 2011), and Western clawed frogs, *Xenopus tropicalis* (microarray: Yanai et al., 2011; RNA-seq: Tan et al., 2013)*. D. rerio* expression data at the zygote developmental stage (0.25 hpf) was excluded because of likely abundance of maternal transcripts, and time points after 4 dpf were excluded due to substantial completion of the developmental program (after Kimmel et al., 1995; Yang et al., 2013).

### Preprocessing microarray and RNA-seq data

Affymetrix and Agilent microarray data were imported directly using the R packages *simpleaffy* and *limma*, respectively (Wilson and Miller, 2005; Ritchie et al., 2015). For both microarray platforms, preprocessing consisted of RMA background correction with quantile normalization (Irizarry et al., 2003). This information was automatically attained by *limma* for Agilent data sets. For Affymetrix and Agilent data, probe sets that mapped to multiple genes or no genes at all were excluded from further analysis. All expression values were then transformed to log-scale using the function *log2(x)* (“Log2-transformed”). For RNA-seq data, raw reads were checked for quality using FastQC (Andrews, 2010). All data sets were of good quality with less than 10% adaptor contamination, thus, no trimming was required. After quality control raw reads were pseudoaligned to species-specific reference transcriptomes using Kallisto to produce read counts (Bray et al., 2016). Read counts were transformed to transcripts per million mapped reads (TPMs). The package *biomaRt* was used to assign corresponding ENSEMBL gene ID(s) to each Affymetrix probe set and RNA-seq transcript (Durinck et al., 2009). For microarray data, the signals of all probe sets mapping to the same gene were averaged. For RNA-seq counts mapped to different transcripts of the same gene were summed. Expression of each gene was averaged across biological replicates for developmental time point. These were the expression values used for downstream analysis.

### Clustering of embryonic stages

We used transcriptomic similarities to classify the embryonic stages of each species in an unbiased manner. First, we determined the number of clusters using an elbow plot method. Specifically, for each species and gene expression profiling technology we performed *k*-means clustering using gene expression for all developmental stages. We varied the number of clusters from 1 ≤ *k* ≤ 9 for microarray and 1 ≤ *k* ≤ 9 for RNA-seq and computed the sums of squares error (SSE, or variance within cluster) for all *k*. To determine an appropriate number of clusters, we used a) the “elbow” effect (or determined the *k* at which additional cluster no longer results in a large reduction in SSE) and b) determined the *k* where clustering of groups maintained temporal order of embryogenesis (i.e., no late stages cluster with early rather than other late stages). Second, to generate clusters of embryonic stages for each species, we hierarchically clustered stages of embryogenesis by similarity in gene expression measured as Spearman’s rank correlations. The resulting dendrograms were partitioned into five groups to determine stage clusters. Description of developmental events were obtained from species-specific references including: zebrafish (Kimmel et al., 1995), chicken (Hamburger and Hamilton, 1992), softshell turtle (Tokita and Kuratani, 2001), both *Xenopus* species (Nieuwkoop and Faber, 1994), and mouse (Graham et al., 2015). For clusters containing more than one embryonic stage, an expression mean was used as the representative expression for that cluster for the remaining analyses.

### Ortholog calling

To identify orthologous genes across taxa we used the sequence-based ortholog calling software package OrthoMCL (Li et al., 2003) for microarray data and FastOrtho (Wattam et al., 2013; an implementation of OrthoMCL) for RNA-seq data. Predicted protein sequences of the reference genomes were organized into orthologous gene groups based on sequence similarity. For each reference proteome, protein and corresponding gene ids were grouped as paralogs when sequence similarity was higher among genes within species than between species. To facilitate downstream analysis of expression conservation across taxa, we removed any orthologous gene groups containing paralogs or losses/absences in one or more species. The resulting one-to-one orthologs were used for all downstream analyses (Supplementary Tables 4 and 5, microarray and RNA-seq, respectively). To assess similarity in microarray and RNA-seq comparison, we compared the one-to-one ortholog sets of three species (zebrafish, chicken, and the Western clawed frog).

### Comparing transcriptomes through embryogenesis across species

For both microarray and RNA-seq data, we assessed transcriptomic similarity at early, middle and late phases of embryogenesis across species by calculating the Spearman’s rank correlation for all pairwise comparisons of species for each of the five clusters of embryonic stages. For microarray data, we excluded one frog (*X. laevis*) from the pairwise comparisons to prevent biasing the outcome as a consequence of high correlations in gene expression between the two congeneric anuran species at each cluster of embryonic stages. Due to the high correlations between these two species, similar results were recovered when *X. tropicalis* was removed from the analysis instead of *X. laevis*. We used permutation analysis to assess whether correlations are higher or lower than expected by chance. Specifically, for each species we randomly assigned stages to a cluster maintaining the original number of stages included in the observed cluster and computed the rank correlation for all pairwise species comparison. We conducted 1000 permutations and assessed significance by comparing the observed rank correlation to distribution of rank correlations generated by permutation analysis. Permutation p-values were defined as the percentile of the observed median Spearman’s rho in the distribution of permuted Spearman’s rho values. Because correlations that are higher or lower than expected by chance were both of interest (i.e., a two-tailed test) the both the percentile and 1-percentile were calculated using the empirical cumulative distribution function in R, the lower of the two is reported.

### Characterizing expression conservation of each gene through embryogenesis across vertebrates

To assess conservation of gene expression for each gene at each cluster of embryonic stages and each node, we calculated a difference in expression rank scaled by the divergence time between the groups (Supplementary Fig. S2). At each node of the phylogeny, we characterized patterns of expression conservation across embryogenesis using the R package Clustering of Time Series Gene Expression Data (*ctsGE*; Sharabi-Schwager and Ophir, 2019). Using *ctsGE*, gene conservation scores of each gene were median-scaled and converted into conservation indices. For each gene, at each cluster of embryonic stages, the standardized values indicate the median absolute distance of that gene from its median conservation score. These standardized values were then converted to index values that indicate whether gene expression conservation was above (1), below (−1), or within (0) the cutoff range (+/− 0.7) of the median value at each time step (here cluster of embryonic stages). Each index of expression conservation across embryogenesis was assigned to a conservation pattern based on median transitions across the assigned significance cutoff (+/− 0.7). For early conservation: similarity decreases through embryogenesis; hourglass: similarity increases and then decreases: inverse hourglass: similarity decreases and then increases; late conservation: similarity increases; invariant: similarity does not vary or does not follow other conservation patterns through embryogenesis (Fig. 1A). Indices assignments are provided in Supplementary Table 6.

To examine enrichment of genes in each conservation pattern, we first calculated the proportion of indices in each conservation pattern and defined the expected number of genes as the equivalent proportion of total genes. We determined significance of enrichment/depletion of genes exhibiting each pattern using a permutation analysis. For each gene we first randomized the order of conservation scores across the clusters of embryonic stages. Second, we characterized conservation trajectory using ctsGE and the index assignment rules described above. Finally, for each iteration we calculated enrichment/depletion of genes compared to the random expectation for each conservation pattern. We repeated this permutation 1000 times for each node and gene expression profiling technology and assessed by comparing the observed enrichment/depletion to the null distribution generated by the permutation analysis. Permutation p-values were defined as the probability of obtaining enrichment/depletion of genes at or above/below the observed number in the permutation set as described above.

## Results

### Variation in gene sets and alignment of embryonic stage clusters across species and gene expression profiling technologies

We identified 1626 and 1793 one-to-one orthologs for the microarray and RNA-seq data in our analysis, respectively, consistent with other comparative gene expression studies spanning vertebrates (3044 avian to human orthologs: Pfenning et al., 2014; 1979 orthologs across Peromyscus mice, Microtus voles, Passeroid birds, Dendrobatid frogs, and Ectodini cichlids: Young et al., 2019). Interestingly, one-to-one orthologs from the microarray and RNA-seq data set were largely non-overlapping with a total of 255 overlapping genes (15.7% and 14.2% of each set of orthologs, respectively). Similarly, one-to-one orthologs from microarray and RNA-seq data sets from the same species were largely non-overlapping as well (~15-20%, Supplementary Table 7). Consistency of this overlap across species suggests that microarray and RNA-seq approaches may target distinct features of the transcriptome.

When we clustered the embryonic stages according to their gene expression patterns we found that, for each species and gene expression profiling technology, a *k* = 5 emerged as the number of clusters at which the reduction in within cluster-variance begins to asymptote (Fig. 3). Importantly, the temporal order of stages across embryogenesis was maintained at *k* = 5 clusters; however, inconsistent sampling across species as well as heterochrony across species resulted in some variation in the major events contained in each cluster of embryonic stages across species and between microarray and RNA-seq platforms (Fig. 4). Stages contained in each cluster and a biological description of embryonic events is provided in Fig. 5 and Supplementary Tables 2 and 3 (microarray and RNA-seq, respectively). The number of stages collapsed into each cluster also varied across species and gene expression profiling technology. Because there was no apparent bias between cluster timing in embryogenesis and the number of stages included (Fig. 5), we concluded that *k* = 5 clusters was appropriate for downstream analyses.

**Figure 3.**
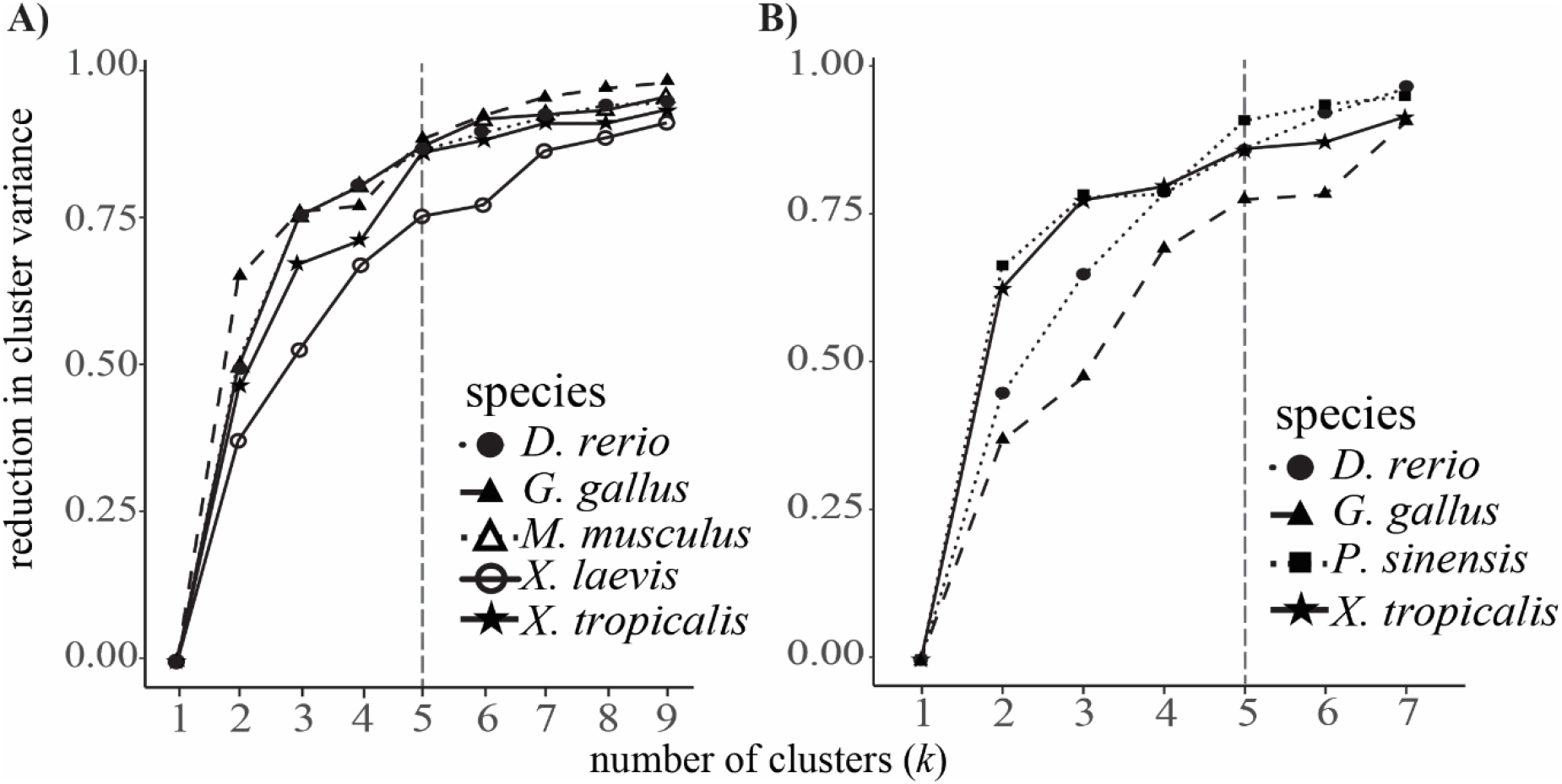
*k*-means clustering of the microarray (A) and RNA-seq (B) data sets analyzed. Reduction of within cluster variance increases as the number of cluster (*k*) increases. Gains asymptote at approximately *k* = 5 (dashed line).

**Figure 4.**
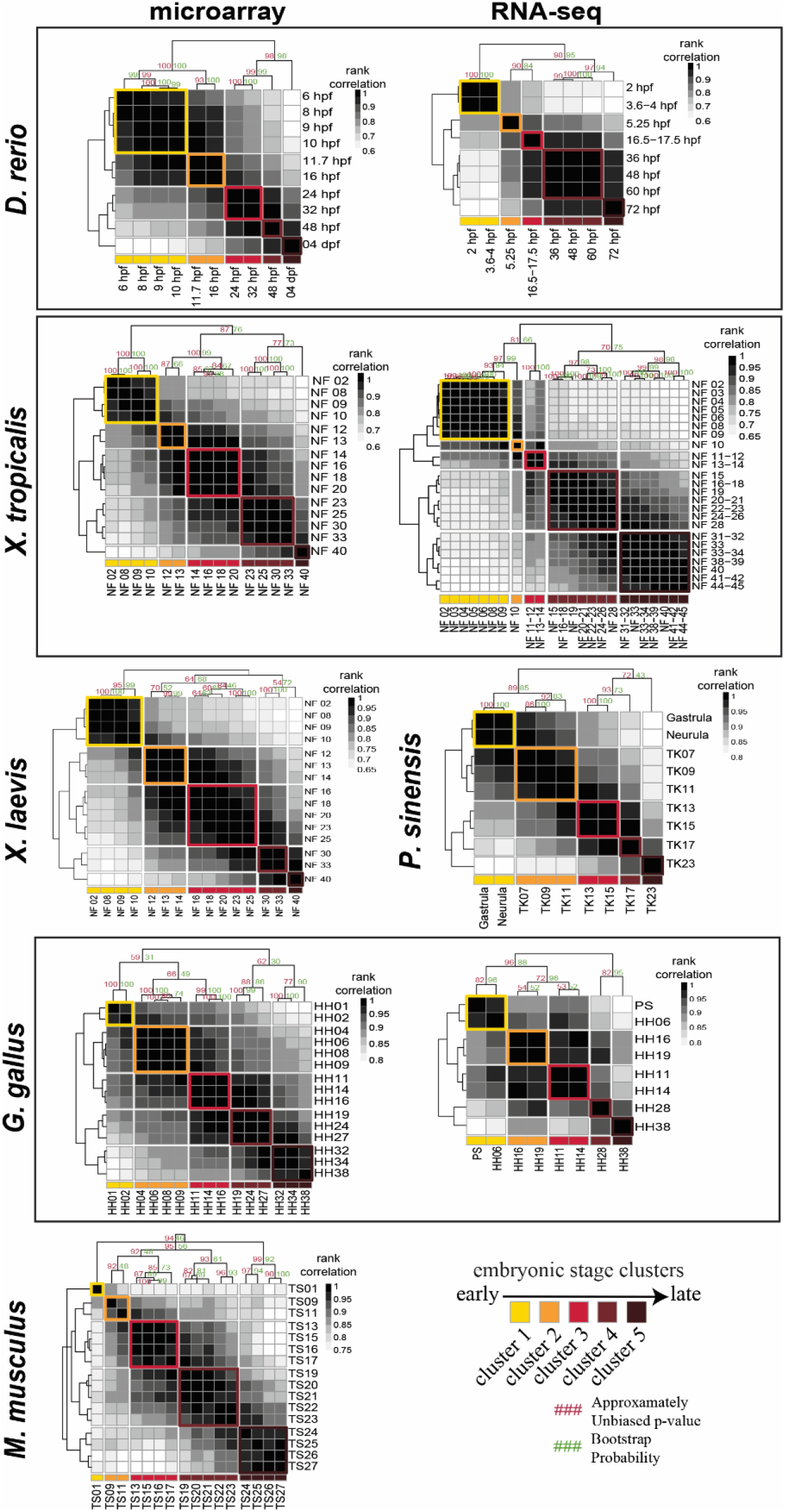
Spearman Rank correlations were used to group stages into five clusters. Shown are all pairwise correlations of stages for all species and both gene expression profiling technologies. Grouping of stages is in indicated color.

**Figure 5.**
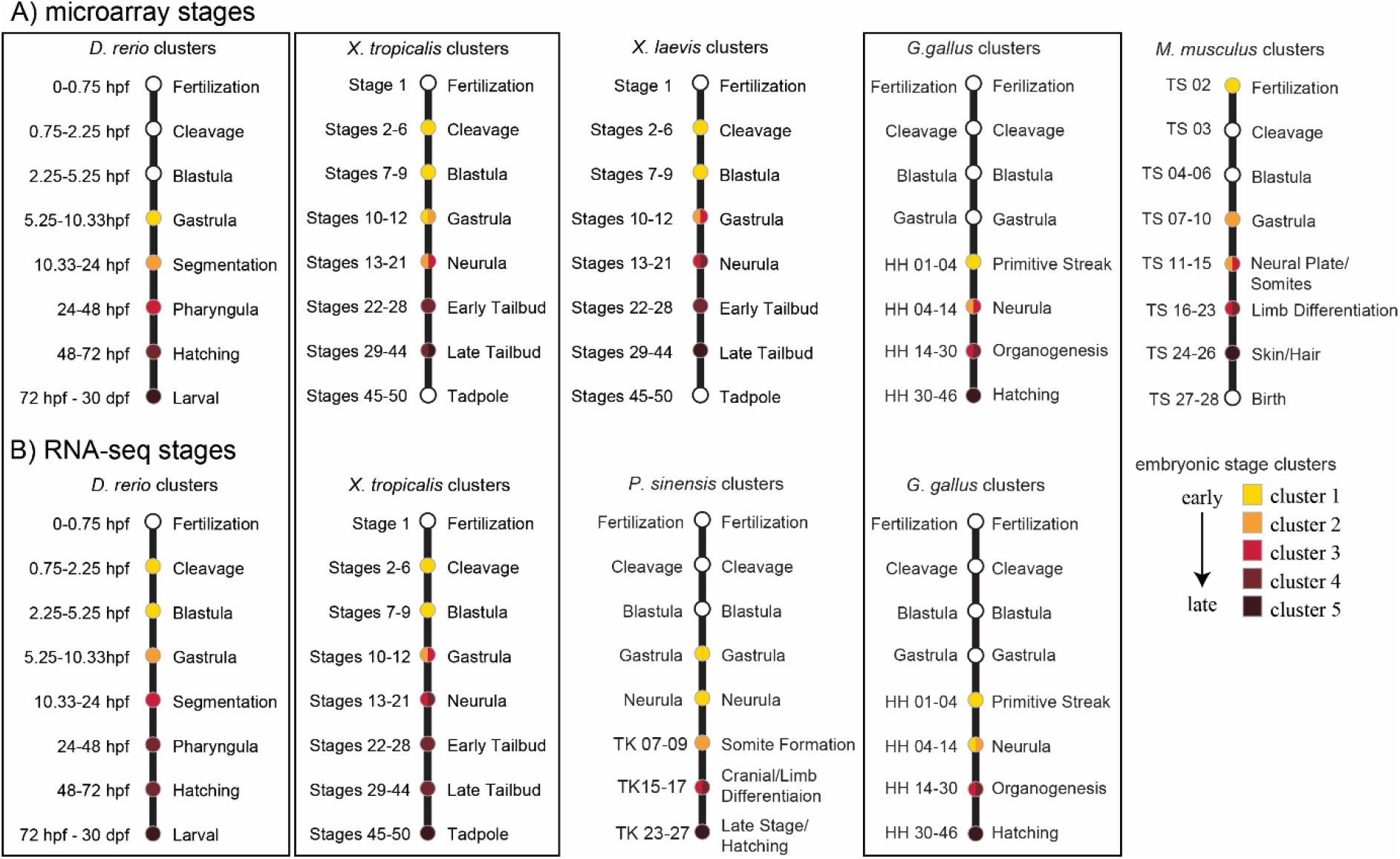
Overlap of embryonic stage cluster and major developmental events. Variation in sampling and heterochrony among species result in differences of developmental events captured by the available data across species and platforms. For three species, gene expression datasets were obtained from both microarray (top) and RNA-seq (bottom) platforms.

### Conservation of gene expression at the transcriptome level

We then quantified interspecific gene expression correlation for each of the *k* = 5 clusters of embryogenesis identified in Fig. 4. Clusters that are correlated across species are more conserved. We found that the pattern of pairwise rank correlations for all one-to-one orthologs (1626 and 1793 for microarray and RNA-seq, respectively) through embryogenesis differed for microarray and RNA-seq comparisons (Fig. 6). In the microarray comparison, median pairwise rank correlations increased mid-embryogenesis, with clusters 2, 3, and 4 having the highest median correlation, and decreased early and late in embryogenesis, with cluster 5 having the lowest median, suggesting a developmental hourglass (Fig. 6A). However, when compared to the null expectation of no gene expression conservation above chance, generated by permuting the cluster assignment of each stage, the observed pattern did not differ from the null (cluster 1: observed median Spearman’s ρ_o_ = 0.46, permutation Spearman’s ρ_p_ = [0.37-0.54], p = 0.43; cluster 2: ρ_o_ = 0.49, ρ_p_ = [0.4-0.53], p = 0.26; cluster 3: ρ_o_ = 0.49, ρ_p_ = [0.42=0.53], p = 0.47; cluster 4: ρ_o_ = 0.48, ρ_p_ = [0.44-0.53], p = 0.35; cluster 5: ρ_o_ = 0.46, ρ_p_ = [0.36-0.52], p = 0.49). In the RNA-seq comparison, median pairwise rank correlations increased to its highest median score at cluster 2 and dropped through later stages of embryogenesis 3, with cluster 5 having the lowest median (Fig. 6B). When compared to the null expectation, median pairwise rank correlation of cluster 2 was significantly higher and correlations of clusters 4 and 5 were significantly lower than expected by chance, providing strong support for a developmental hourglass (cluster 1: ρ_o_ = 0.65, ρ_p_ = [0.56-0.68], p = 0.24; cluster 2: ρ_o_ = 0.67, ρ_p_ = [0.52-0.69], p = 0.005; cluster 3: ρ_o_ = 0.62, ρ_p_ = [0.51-0.68], p = 0.49; cluster 4: ρ_o_ = 0.57, ρ_p_ = [0.57-0.68], p = 0.003; cluster 5: ρ_o_ = 0.53, ρ_p_ = [0.59-0.69], p = 0.008).

**Figure 6.**
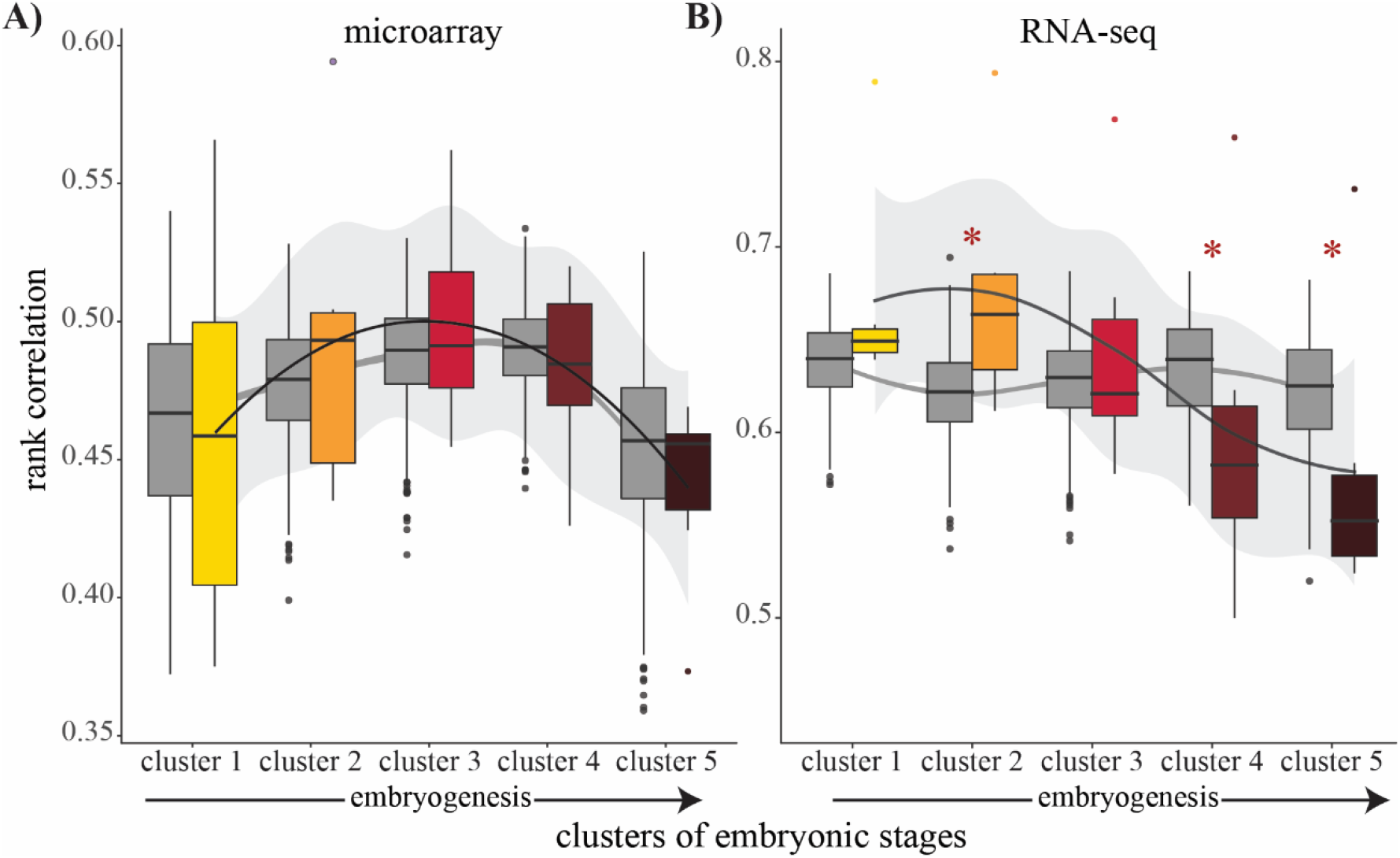
Spearman rank correlations for pairwise comparisons of species at each cluster of embryogenesis for microarray (A) and RNA-seq data (B). Gene expression correlations (as a measure of conservation) vary through embryogenesis for both microarray and RNA-seq data and appear to show support for a developmental hourglass. Colored boxes indicate observed correlations; grey boxes indicate rank correlations after permutation analysis randomizing stage association with cluster. Asterisks indicate that the observed median correlation differs significantly from the null expectation at *p < 0.01*. Note that only the RNA-seq data showed a temporal pattern (consistent with the early conservation hypothesis) that differed from the null expectation (i.e., no temporal pattern present).

### Enrichment and overlap of expression conservation patterns across the phylogeny

Next, we used ctsGE time series analysis to characterize patterns of gene expression conservation across embryogenesis at each phylogenetic node. For the microarray data set, gene conservation scores yielded a total of 90, 90, 87, 89 patterns of expression conservation in anurans, amniotes, tetrapods, and vertebrates, respectively. For the RNA-seq data set, gene conservation scores yielded a total of 90, 51, and 49 patterns in amniotes, tetrapods, and vertebrates. We calculated the expected number of genes for each pattern as equivalent to the proportion of total patterns (Fig. 7, treemap insets; Tennekes, 2017). Across vertebrates, with the exception of inverse hourglass in the RNA-seq analysis and time-invariance in both platforms, all patterns were exhibited by significantly more genes than expected by chance (Fig. 7). This enrichment of genes across all conservation patterns and the depletion of time-invariant genes deviated significantly from the null expectation (microarray: early conservation p = 0, hourglass p = 0, inverse hourglass p = 0, late conservation p = 0, time invariance p = 0; RNA-seq: early conservation p = 0, hourglass p = 0.02, inverse hourglass p = 0.4, late conservation p = 0, time invariance p = 0) Enrichment/depletion of genes for each conservation pattern across anurans (microarray only), amniotes, and tetrapods are provided in Supplementary Fig. S3. p = 0 indicates that none of the 1000 permutation iterations resulted in enrichment/depletion at or above/below the observed level.

**Figure 7.**
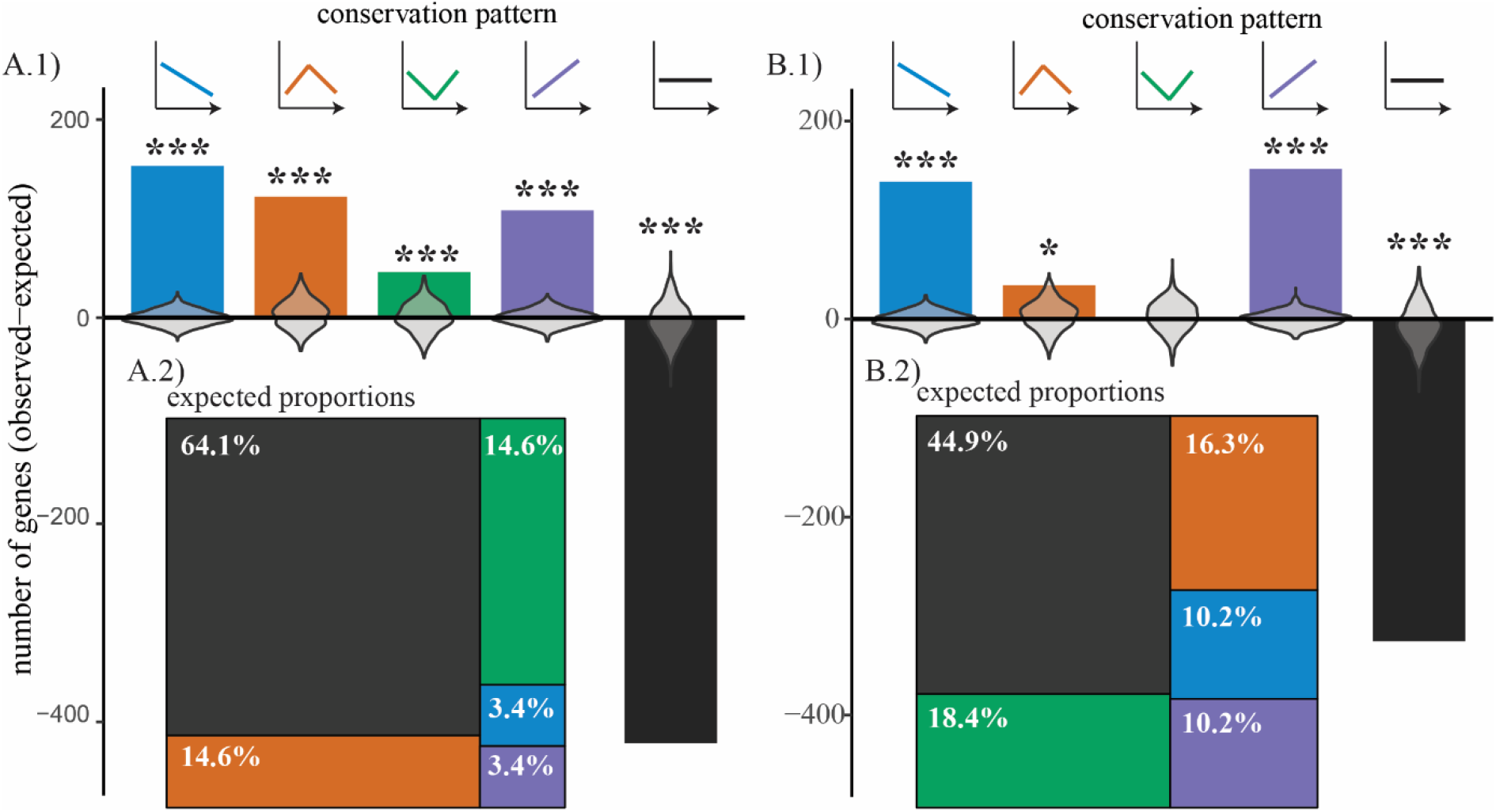
Enrichment of patterned gene expression conservation across vertebrates. The expected number of genes for a given temporal expression pattern is determined by the proportion of trajectories of each conservation pattern (A.2: microarray; B.2: RNA-seq). The proportion of trajectories differed across conservation patterns and between platforms (A.2 and B.2). The enrichment/depletion values from permutation analysis (1000 iterations) are shown as violin plots (A.1 and B.1). The number of genes associated with all conservation patterns were enriched compared to random expectation, except for the inverse hourglass (RNA-seq only), whereas the number of genes with no relationship between expression conservation across embryogenesis (A.1: microarray; B.1: RNA-seq) was significantly less than expected by chance. For all plots colors indicate the expression conservation pattern. Enrichment at other evolutionary nodes (tetrapods, amniotes, and anurans) are provided in Supplementary Fig. S3.

Finally, we compared conservation pattern assignment for the 255 one-to-one orthologous gene groups shared between microarray and RNA-seq gene sets and found that across platforms gene expression patterns are not concordant at the major phylogenetic nodes. Specifically, only 25% of genes (63/255) in amniotes, 19% (49/255) in tetrapods, and 20% (52/255) in vertebrates exhibited concordant expression conservation patterns.

## Discussion

In the present study, we examined patterns of expression diversification throughout embryogenesis in six vertebrate species using a comparative analysis of 112 microarray and 74 RNA-seq data sets (Fig. 1B). First, using an unbiased approach, we clustered stages of embryogenesis within species and compared expression conservation of those clusters across species (Figs. 3-5; Supplementary Tables 2 and 3). This approach allowed for the inclusion of all available stages of embryogenesis and removed bias that could result from comparing only selected stages. Second, we used a permutation analysis to generate a null expectation. We observed conservation estimates against this null expectation to characterize transcriptome-level diversification through embryogenesis across species (Fig. 4). Finally, we characterized the expression conservation of each gene at each node of the phylogeny (Fig. 1B; Supplementary Fig. S2) to examine how expression conservation patterns vary through evolutionary time (Fig. 6).

Over the past decade, the debate of whether diversification of embryogenesis follows generalizable rules has been reinvigorated by the ability to test predictions of the hourglass, developmental burden, and other hypotheses (Fig. 1A) on a genomic scale. Enabled by the increase in ‘omics-level data and next generation sequencing accessibility, a number of studies have by now explored patterns of diversification in gene expression though embryogenesis across species (Fig. 2). These studies have found mixed support for the hourglass and other models of diversification across species (Fig. 2B-D; reviewed in: Irie, 2017; Liu and Robinson-Rechavi, 2018). Differences among studies could reflect differences in species compared as some studies span phyla (de Mendoza et al., 2013; Levin et al., 2016; Hu et al., 2017) and others are restricted to internal nodes of the vertebrate phylogeny (e.g., amniotes; Wang et al., 2013). However, our quantitative literature analysis did not indicate an effect of divergence time on the characterization of an hourglass model of divergence (Fig. 2C). Alternatively, selection of developmental stages or specific gene sets to compare may lead to different results and interpretations. Finally, studies that compare gene expression similarity at the transcriptome level rarely test against a null hypothesis (Young and Hofmann, 2019). Such a test is critical because the degree of variation expected through developmental stages across species is unknown (Church and Extavour, 2020).

Aligning stages of early animal development is complicated by taxon-specific sampling practices as well as pervasive heterochrony in developmental events across distantly related species, leading some researchers to question the validity of anatomical hourglass hypotheses (Bininda-Emonds et al., 2003). We found that while expression varies in similar ways through embryogenesis across species, stages of embryogenesis did not always consistently cluster together within species (Fig.3 and 4; Supplementary Tables 2 and 3). These differences likely reflect both biological variation in the molecular timing of developmental events, technical variation in sampling procedures across species, and a lack of available data sets particularly at early embryonic stages. Because clustering has the advantage of being unbiased, and no systematic bias in sampling was apparent, we moved forward by comparing gene expression at each embryonic stage cluster across species. Future studies with systematic sampling of embryogenesis across species could disentangle the source (biological or technical) of variation in stage clustering across species.

Consistent with the mixed support for the hourglass and other models of developmental divergence (e.g., early conservation or inverse hourglass) found across studies, our comparisons of expression variation in all one-to-one orthologs present across species in the microarray and RNA-seq data yielded significant but different patterns. Specifically, the observed gene expression correlations (as a measure of conservation) differed through embryogenesis for both microarray and RNA-seq, yet only the RNA-seq data showed a pattern that differed significantly from the null expectation that gene expression levels should be invariant of developmental time (Fig. 6). For the RNA-seq datasets, we found a significant increase in expression correlation over the null expectation in developmental time cluster 2 followed by a significant reduction in expression correlation in the later clusters 4 and 5. Though clusters 1 and 2 display similar gene expression correlations, suggesting an early conservation pattern, cluster 1 does not significantly differ from the null expectation. Inconsistencies between the microarray and RNA-seq data sets could result from differences in taxon sampling (Fig. 1B), the embryonic stages and resulting clusters that were included (Figs. 4 and 5), and/or systematic differences in which aspects of the transcriptome were captured by these distinct platforms. In fact, transcriptome-level gene expression profiles quantified using RNA-seq and microarray technologies have been shown to be correlated especially for highly expressed genes, but variation between technologies is also commonly reported (Marioni et al., 2008; Malone and Oliver, 2011; Trost et al., 2015). Whether these differences reflect superiority of one technology over the other is unclear. Instead technical differences between the two approaches may capture different elements of the transcriptome, in which case the two approaches should be viewed as complementary (Kogenaru et al., 2012). In our data sets, we found little overlap of one-to-one orthologs between microarray and RNA-seq, with only ~15-20% of the genes contained in both analyses. Further, of those one-to-one orthologs contained in both microarray and RNA-seq data sets, only ~20% shared conservation pattern assignments (Fig. 8). This illustrates a major challenge for comparative analyses, like our present study, that aim to capitalize on the vast amounts of publicly available transcriptome datasets to test biological hypotheses.

**Figure 8.**
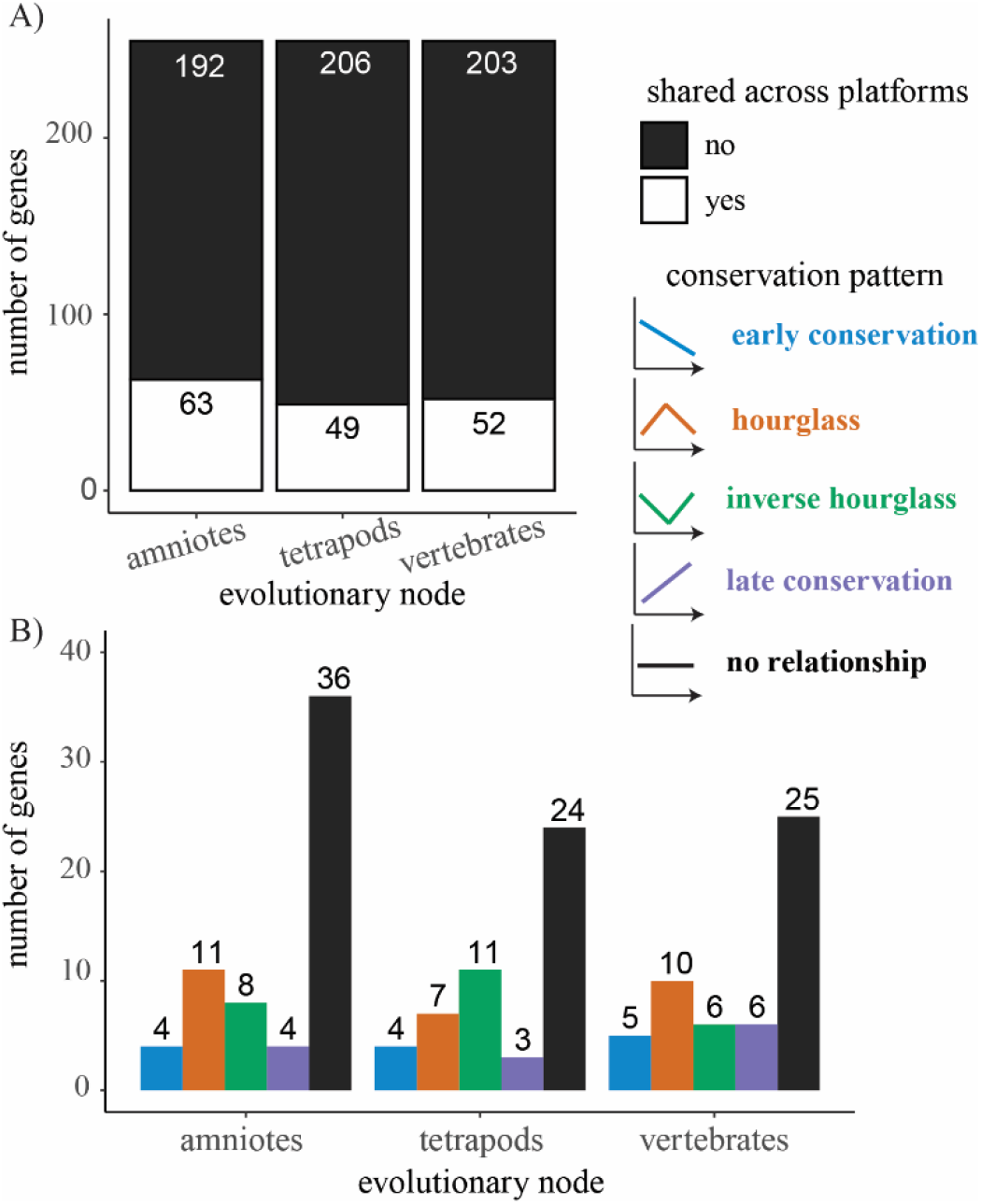
255 one-to-one orthologs shared between microarray and RNA-seq data sets largely differed in expression conservation pattern (A). Most genes with shared expression conservation patterns were “no relationship” genes followed by hourglass or inverse hourglass genes (B).

Inconsistency across studies and gene expression profiling technology also suggests that comparing the whole transcriptome may not be appropriate. Specifically, patterns of expression conservation at the whole transcriptome-level may be biased by abundant and constitutively expressed genes (e.g., see Piasecka et al., 2013). Whole genome approaches have the great advantage of being unbiased, which allows for the identification of novel gene associations with phenotypes and/or gene interactions that would be missed using a candidate gene approach. However, we know that variation is not equivalent across levels of biological organization. For example, studies comparing mRNA and protein levels have found only modest overall correlation (Foss et al., 2007; Fu et al., 2009; Ghazalpour et al., 2011; Vogel and Marcotte, 2012). Further, studies of character homology have shown that even anatomically and physiologically similar homologies can differ in underlying developmental and molecular mechanisms (Wagner, 1989; Wilkins, 2002; McGary et al., 2010; Young and Wagner, 2011). To address this issue, we also used a time series analysis to characterize the conservation pattern of each gene at each node of the phylogeny (Fig. 7), in addition to a transcriptome-level comparisons through embryogenesis across species. Across vertebrates, we observed a significant enrichment of all patterned conservation models (early conservation, hourglass, inverse hourglass, and late conservation) above the expected number and a large depletion of genes whose expression was invariant of developmental time (Fig. 7; Supplementary Fig. S3) in both microarray and RNA-seq data sets.

Both the enrichment of genes exhibiting different patterns and the proportion of overall genes with distinct patterns varied across evolutionary nodes (e.g., tetrapods, amniotes, and anurans, Fig. 1B) and between gene expression profiling technologies (Supplementary Fig. S3). These results suggest that divergence in gene expression through embryogenesis may depend on the evolutionary distance covered by any given analysis. Follow-up studies, including phylogenetic comparative analyses of both closely and distantly related species are needed to better understand these patterns and their implications for generating biological diversity. In addition, time series analyses that characterize expression divergence of individual genes or gene sets should be used to test hypothetical mechanisms of constraint. For example, we might ask whether regulatory interactions or temporal and spatial expression patterns of a gene follow a correlated dynamic pattern through embryogenesis.

## Conclusions

Taken together, our results provided strong support for a patterned embryonic gene expression diversification across vertebrate species. However, the gene groups and evolutionary nodes under which each pattern emerges remain unknown. By combining unbiased clustering of embryonic stages and explicit tests against a null hypothesis our research demonstrates a critical need for broad evolutionary sampling and systematic examination of developmental stages across species to better characterize gene expression diversification in embryogenesis.

## Acknowledgements

We thank D. Wylie for consulting on data analysis, S. Campbell for helping with quantitative literature analyses, and, S. Caro, C. Friesen, I. Miller-Crews, M. Rodriguez Santiago, and T. Solomon-Lane Santiago for discussions that improved this manuscript. This work was funded in part by a University of Texas at Austin Undergraduate Research Fellowship to P.S.B. and a grant from the NSF BEACON Center for the Study of Evolution in Action, DBI-0939454 to H.J.G, A.H., H.A.H., and R.L.Y.

## Supplementary Tables and Figures

Supplementary Table 1. Studies and data included in literature analyses Fig. 2.

Supplementary Table 2. Description and clustering of embryonic stages in species with microarray data

Supplementary Table 3. Description and clustering of embryonic stages in species with RNA-seq data

Supplementary Table 4. Orthologous gene groups for microarray datasets

Supplementary Table 5. Orthologous gene groups for RNA-seq datasets

Supplementary Table 6. Gene conservations pattern indices

**Supplementary Table 7.** Number and percentage of overlapping one-to-one orthologs in microarray and RNA-seq datasets

| Species | # of overlapping orthologs | % of orthologs (microarray) | % of orthologs (RNA-seq) |
| --- | --- | --- | --- |
| *Gallus gallus* | 264 | 18.7% | 18.0% |
| *Xenopus tropicalis* | 345 | 21.2% | 19.2% |
| *Danio rerio* | 335 | 20.6% | 16.2% |

**Supplementary Figure S1.**
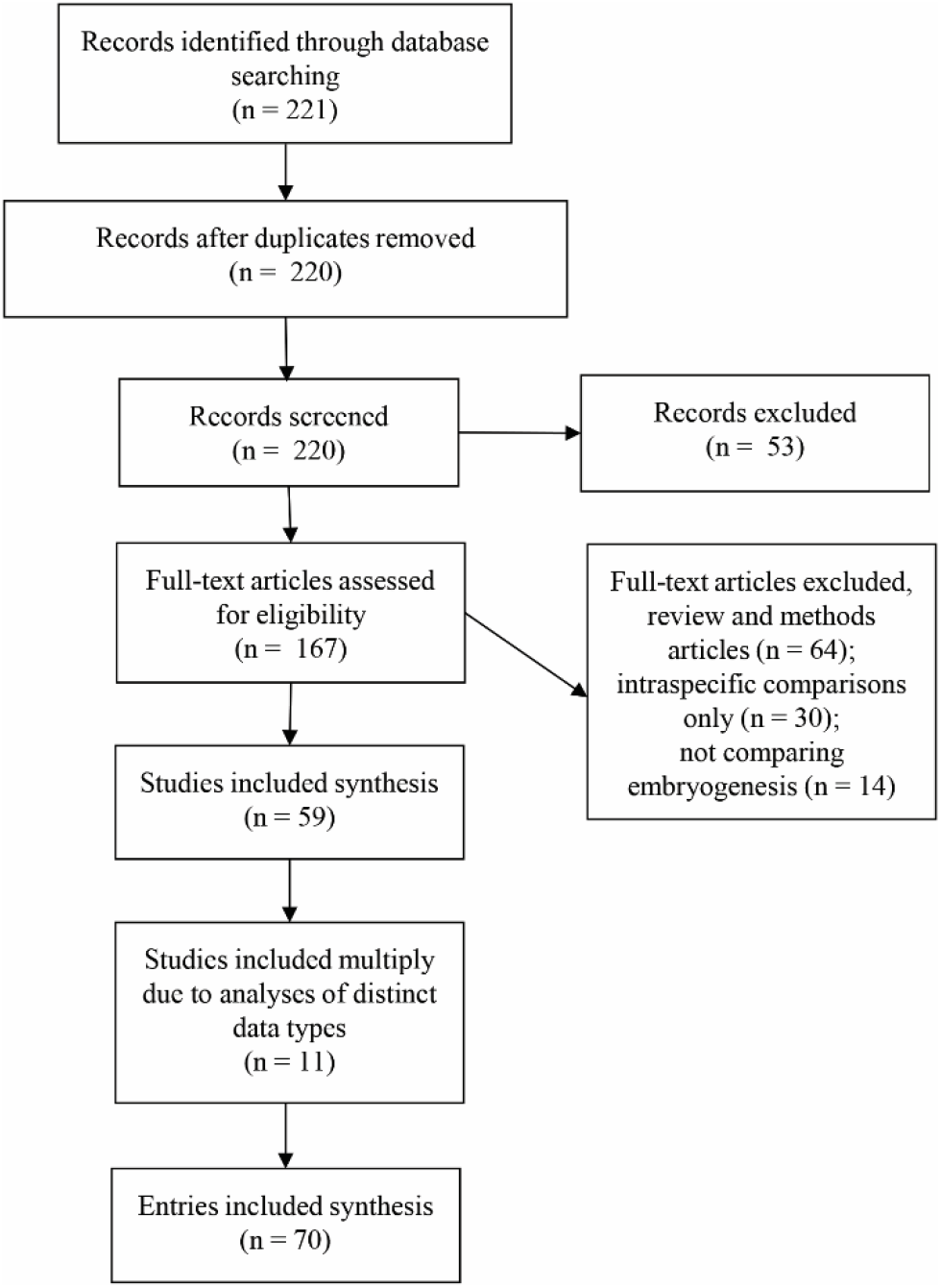
Modified PRIMSA (Moher et al., 2009) flow chart. 220 unique articles retrieved from Web of Science using the following search criteria topic (TS) equals “phylotypic” OR “developmental AND hourglass” on March 25, 2020. We removed 53 articles that were not about the developmental hourglass hypothesis (e.g., in microbiology, “phylotypic” is sometimes used to classify a group of organisms by their phenetic relationship). We removed an additional 64 methods and review articles that did not include data analysis, 30 articles that included intraspecific comparisons only, and 14 articles that compared development of individual phenotypes (e.g., craniofacial morphology, brain, limbs, and heart) rather than embryogenesis. For each of the remaining 59 unique articles (Supplementary Table 1), the pattern of variation (early conservation, hourglass, inverse hourglass, late conservation, or no relationship) was reported for the type of data analyzed (gene expression, molecular evolution, morphological, or regulatory). 11 studies included analyses of data from different biological levels of organization (e.g., molecular evolution and transcriptomics). These analyses were treated as independent resulting in a total of 70 entries. Finally, four studies compared species with unresolved divergence times. These articles were excluded from divergence time comparisons in Fig. 2C. Divergence time estimates were obtained from TimeTree (Kumar et al., 2017). A linear regression model was used to assess increase in research focus and gene expression-based studies and test whether an increased proportion of studies find an hourglass.

**Supplementary Figure S2.**
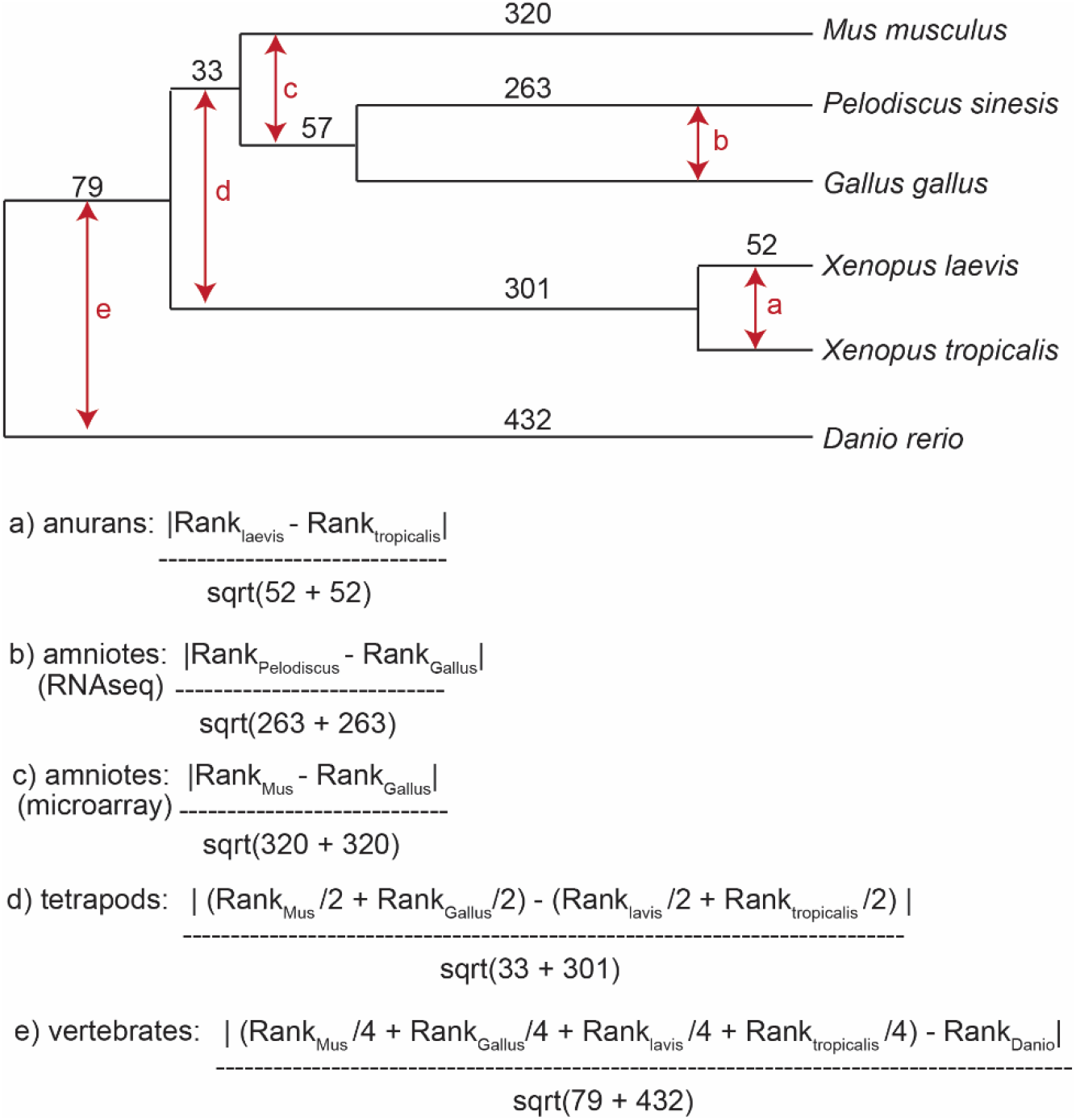
To characterize expression conservation of each gene at each embryonic stage cluster at each node of the phylogeny we calculated the similarity in rank expression scaled by the divergence times of the included species.

**Supplementary Figure S3.**
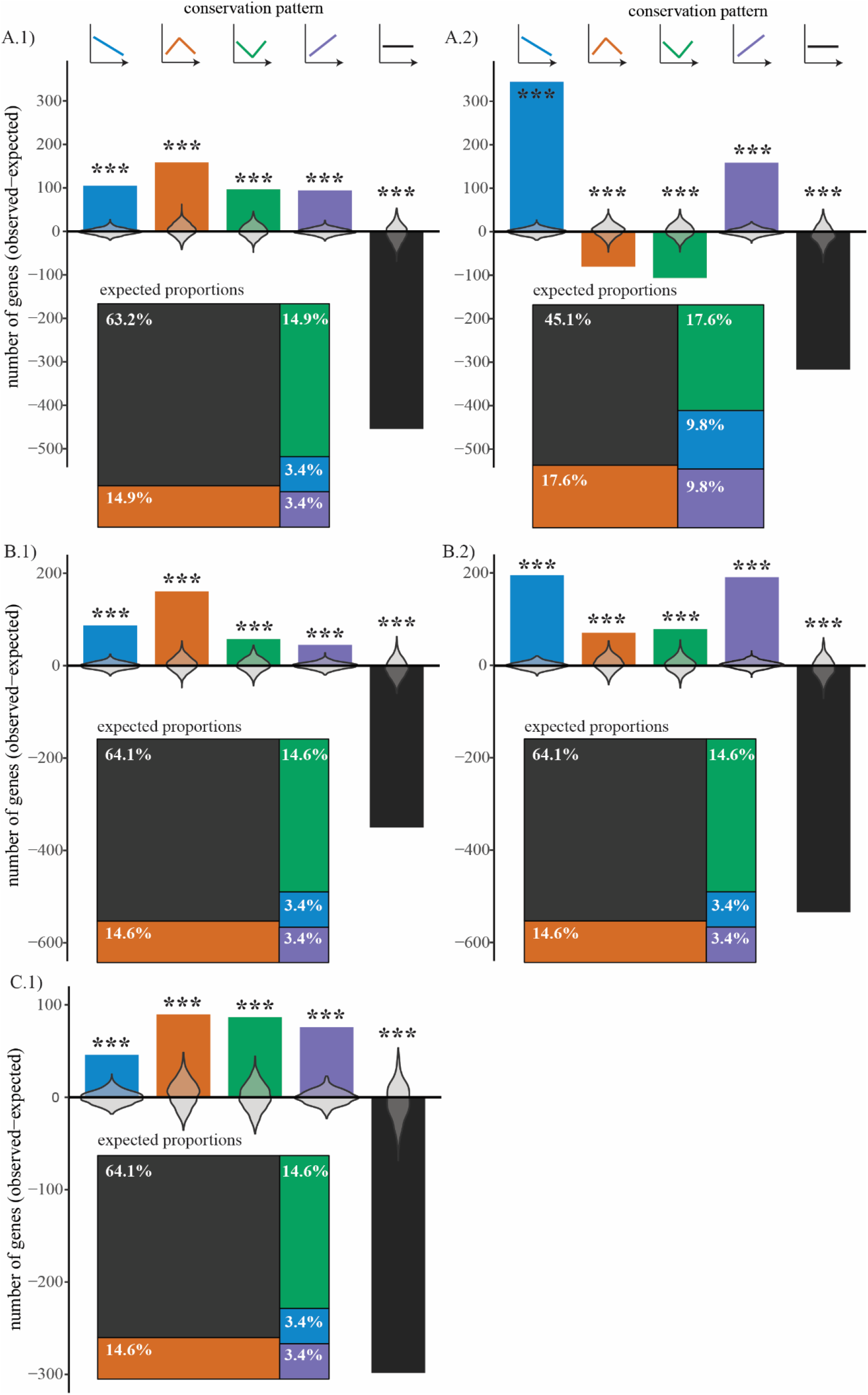
Enrichment or depletion of gene expression conservation patterns at available evolutionary nodes amniotes (A), tetrapods (B), and anurans (C) for microarray (1: left) and RNA-seq (2: right). Proportion of patterns determines the expected proportion of genes for each conservation pattern and are shown as associated treemap plots (Tennekes, 2017). The enrichment/depletion values from permutation analysis (1000 iterations) are shown as overlaying violin plots. For all plots colors indicate the expression conservation pattern. *** indicate significance at p = 0.

